# Opposing influences of sensory and response history in larval zebrafish

**DOI:** 10.64898/2026.07.30.741669

**Authors:** Ashrit Mangalwedhekar, Sydney Hunt, Iacopo Hachen, Armin Bahl

## Abstract

Variability is a prominent feature of animal behavior across species. Despite highly controlled experimental conditions, the same individual often responds differently to repeated identical stimulation. Part of this variability can be explained by the sequence of previous sensory stimuli and decision-making events – the trial history. Most studies about trial history are limited to animals that have a cortex, such as rodents or primates, observed as they perform learned cognitive tasks. It is currently unknown whether trial history can shape the behavior of animals lacking cortical structures during untrained, naturalistic behaviors. Here, we address this question in larval zebrafish, performing the optomotor response, an innate sensorimotor behavior in which animals turn in the direction of whole-field visual motion. We observed that a substantial proportion of variability can be explained by recent stimulation and decision-making events, with influences lasting for tens of minutes. Intriguingly, sensory and response histories bias the current response in opposite directions – repulsive and attractive, respectively – analogous to what has been previously reported in humans and rodents. An integrator model operating across multiple timescales explains a considerable fraction of response variability based on trial history alone. Our findings demonstrate that history dependency in animal behavior is not an exclusive feature of higher-order cortical computation after learning, but that it reflects a fundamental and evolutionarily shared property of vertebrate sensorimotor systems.

**SIGNIFICANCE STATEMENT:** Behavioral variability is often attributed to noise in sensory and neural processing. We show instead that much of this variability can arise predictably from an animal’s recent sensory and response history. In freely swimming larval zebrafish performing an innate visual behavior, previous stimuli and responses exert opposing, long-lasting influences: sensory history biases behavior away from the past, whereas response history promotes repetition. Such trial-to-trial effects are increasingly recognized in cognitive neuroscience but, to our knowledge, have not previously been demonstrated in a nonmammalian vertebrate. Their presence in zebrafish, a model system offering brain-wide neural access and powerful molecular tools, creates new opportunities to identify potentially conserved circuit mechanisms underlying persistent sensorimotor computations.

**HIGHLIGHTS:**

- Visual motion and response histories have opposing influences on behavior
- Stimulus history negatively biases response direction over tens of seconds
- Animals tend to repeat responses across trials, a positive bias that slowly increases over minutes
- A deterministic model applied to experimental sessions explains considerable fractions of response variability

## INTRODUCTION

Animal behavior is inherently variable: seemingly identical sensory inputs often generate different responses. Even within the same individual, repeated stimulus presentations can lead to different outcomes across trials. A part of this variability may simply arise from noisy sensorimotor transformations in the nervous system. However, the influence of previous stimuli and motor decision events plays an important role as well, but it is often neglected when analyzing behavior and exploring neural dynamics. Several studies have reported that previous stimuli, decision events, and associated rewards influence the response in the current trial. Such hysteresis in perceptual decision-making tasks has been well characterized in rodents and primates, including humans (Abrahamyan *et al*., 2016; Akrami *et al*., 2018; Lak *et al*., 2020; Hachen *et al*., 2021).

Experiments in perceptual decision-making often involve a trained subject responding to a well-defined or random series of sensory stimuli. Trials are composed of stimulus presentation periods and decision event recordings, with or without reward contingencies. Each of these components may influence the current response differently. For example, in humans, it has been reported that, despite random stimulus order, perception is biased towards stimuli in the recent past (Fischer and Whitney, 2014). Contrary to this finding, other works have reported that past stimuli push the current decision away from the previous ones – often referred to as repulsion in psychophysics literature – whereas previous decisions pull the current response towards the same one – often referred to as attraction, repetition, or serial dependence (Fritsche, Mostert and Lange, 2017). Similar effects have been reported in mice, rats, monkeys, and humans (Akaishi *et al*., 2014; Abrahamyan *et al*., 2016; Busse *et al*., 2017; Urai, Braun and Donner, 2017; Akrami *et al*., 2018; Urai *et al*., 2019; Lak *et al*., 2020; Hachen *et al*., 2021). While most studies use visual stimuli, some works have reported qualitatively identical effects using tactile, auditory, and olfactory tasks (Lak *et al*., 2020; Hachen *et al*., 2021). A consistent finding across all these studies is that even when sensory inputs are identical, history-dependent biases can alter responses and introduce considerable variability in behavior.

Recent work in the larval zebrafish suggests that stimulus-dependent history and attention-like processes contribute to behavioral variability (Krishnan *et al*., 2025; Tanaka and Portugues, 2025; Zhao *et al*., 2026). In larval zebrafish, sensorimotor decision-making processes are often studied through the innately present optomotor response. When presented with whole-field visual motion, larval zebrafish turn and swim in the direction of perceived motion (Orger *et al*., 2000). Using this paradigm, studies have shown that larval zebrafish do not solely rely on momentary information but integrate motion information from seconds to tens of seconds to modulate behavior (Mu *et al*., 2019; Bahl and Engert, 2020; Dragomir, Štih and Portugues, 2020; Markov *et al*., 2021; Yang *et al*., 2022; Tanaka and Portugues, 2025). In these studies, larvae are presented with a sequence of stimuli of random order. Responses are then averaged to build computational models of how larvae respond to these stimuli. However, the influence of the trial sequence remains unexplored. Additionally, some of these studies only focus on forward motion in head-fixed or fictively swimming electrophysiology configurations, leaving open questions on how history dependencies may influence turning behavior in freely swimming animals (Mu *et al*., 2019; Markov *et al*., 2021; Yang *et al*., 2022; Tanaka and Portugues, 2025; Zhao *et al*., 2026).

Our objective in the present study is to systematically dissect history-dependent biases and quantify their influence on sensorimotor decision-making through analyses of freely swimming behavior in the context of the optomotor response. Specifically, we seek to explain to what extent behavioral variability arises from preceding trials and motor outputs. First, we show that the optomotor response is variable with strong trial-to-trial history-dependencies, even in stimuli with minimal sensory noise. Second, using motion stimuli of varying degrees of uncertainty, we find that the sensory and response histories have opposing effects on the current response. The tendency to repeat a response increases over experimental time, which consequently reduces optomotor performance. Third, we propose a multiple integrator model that captures behavioral dynamics across three timescales: seconds (within-trial), tens of seconds (trial-to-trial), and hours (across the experiment). Thus, our experimental analyses and joint modeling suggest that behavioral trial-to-trial variability in sensorimotor tasks can largely be explained by deterministic history-dependent features, without requiring explicit modeling of intrinsic perceptual or nervous system noise.

## RESULTS

### Optomotor behavior is modulated by trial-to-trial history

Sensorimotor decision-making is usually studied by averaging responses across randomly ordered trial repeats. However, behavior may considerably depend on both the recent stimulation and response history. To investigate such effects, we focused on the optomotor behavior of larval zebrafish that swam freely in circular arenas (**Fig. 1a**) and were stimulated from below with sinusoidal gratings or random-dot motion (**Fig. 1b** and **Methods**). Using real-time tracking of body position and orientation, we updated stimuli such that the direction of motion always remained perpendicular to the orientation of the larva (Bahl and Engert, 2020; Dragomir, Štih and Portugues, 2020; Slangewal *et al*., 2026). This configuration enables continuous presentation of the same stimulus from the perspective of the fish. Swim bout events occurred approximately once per second and were detected online (**Fig. 1c**). We repeatedly presented larvae with motion stimuli for over two hours (**Fig. 1d**). Every trial consisted of 30 seconds of continuous whole-field motion, interleaved by an inter-trial interval of 40 seconds without motion (**Methods**). Fish responded with directed turning bouts with an interbout interval of around 0.5 s for both stimuli (**Fig. S1a,b,f,g**). We assigned a single response value *R_n_* to each trial by averaging all turning angles during visual motion, followed by binarization into right (R) and left (L) response trials. (**Fig. 1d**, right). Such a binarized trial response was labeled as “correct” if it went in the same direction as the stimulus (**Fig. 1d**, right; first and third example trials) or as “incorrect” otherwise (**Fig. 1d**, right; second example trial).

**Figure 1.**
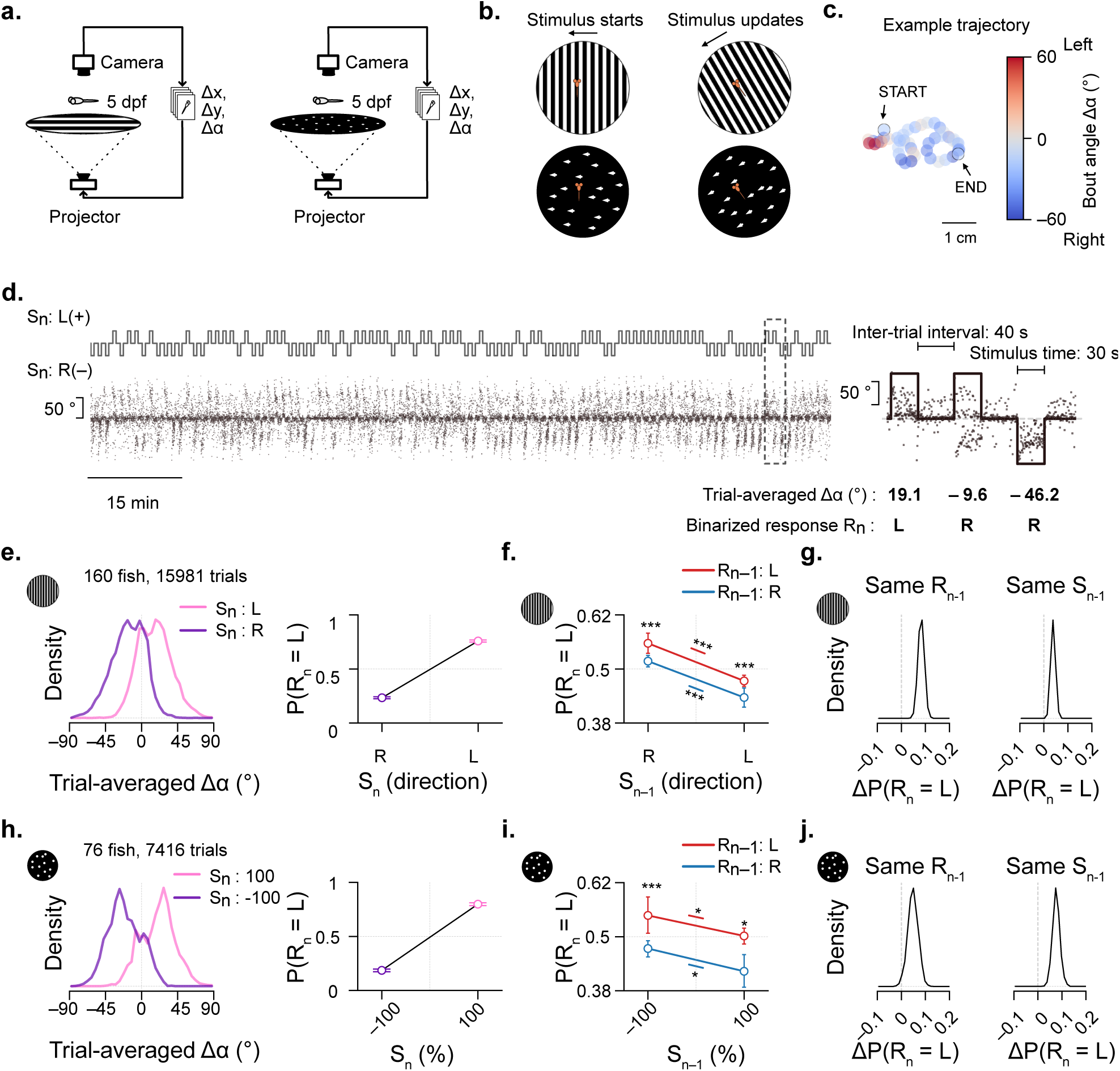
Trial-to-trial history dependency of the optomotor response. **a**, Schematic of the behavioral setup and experiments with the moving grating stimulus (left) and the 100% coherence random-dot-motion stimulus (right). **b**, Schematic representation of fish-locked stimulus. The direction of motion is locked to the body orientation of the fish in real-time during the stimulus period (above: moving gratings, below: random-dot motion). **c**, Trajectory of one example trial. Swim bouts are illustrated by solid colored dots. **d**, Schematic representation of stimulation sequence order (top) and respective swim bout turn angle (gray scatter below) of one example fish. Each dot represents one swim bout. Stimuli and bouts of three example trials are shown on the right. Binarized trial responses (left, L; right, R) are based on averaging all swim bouts within each trial. **e,h**, Left: Distribution of trial-averaged turn angles for the grating (e) and 100% coherence random-dot-motion experiments (h). Right: Probability of a left response (binarized trial-averaged turn angle) for right and the left-ward moving gratings (e) and the 100% coherence random-dot-motion (h) stimuli. **f,i**, Probability of a left response, conditioned on the previous stimulus direction (*S_n_*_−1_) and on the previous response *R_n_*_−1_ (color-coded). For all response condition comparisons within the same previous stimulus (vertical): ***p < 0.001 (bootstrap test). For comparison between previous stimulus directions within response directions (horizontal): ***p < 0.001 for the grating (e) and *p < 0.05 for the 100% coherence random-dot-motion stimulus (i). **g,j**, Differences in response probabilities are derived from (f,i), for gratings (g) and random-dot motion (j). The *S_n_*_−1_ effect is defined as *ΔP* = (*R_n_* = *L*; *S_n_*_−1_ = *L*) − *P*(*R_n_* = *L*; *S_n_*_−1_ = *R*), with merged data from *R_n_*_−1_ = *L* and *R_n_*_−1_ = *R*. The *R_n_*_−1_ effect is related to *ΔP* = *P*(*R_n_* = *L*; *R_n_*_−1_ = *L*) − *P*(*R_n_* = *L*; *R_n_*_−1_ = *R*), with merged data from *S_n_*_−1_ = *L* and *S_n_*_−1_ = *R*. Statistics for the *S_n_*_−1_ and *R_n_*_−1_ effects, comparing grating and 100% coherence random-dot-motion stimuli, were computed by a permutation test (**Methods**). p = 0.16 for the *S_n_*_−1_ effect and p < 0.05 for the *R_n_*_−1_ effect. N values (number of fish and trials) for both stimuli are given in (e,h). All bootstrap tests have been performed by merging trials from all fish (**Methods**). See also **Fig. S1**.

We first sought to quantify history effects in a regime where stimuli are strong (**Fig. 1e–j**), i.e., for strong contrast sine gratings (**Fig. 1e–g**) and 100% coherence random-dot-motion (**Fig. 1h–j**, **Methods**). For both stimuli, we found that the mean trial-averaged turn angles vary considerably across trials **(Fig. 1e,h**, left), with around a quarter of the trials showing negative values where animals, on average, swam against the motion stimulus. The binarized responses corroborated this feature (**Fig. 1e,h**, right). The proportion of incorrect trials indicates that, despite the strong motion stimuli, motor responses are not always aligned with the direction of the visual stimulus.

Could these variations arise from the influence of previous trials? Each trial is associated with two components: the stimulus direction (*S*) and the larva’s response (*R*). To disentangle the effect of each of these components, we quantified the average response in a given trial n (*R_n_*), conditioned on the previous stimulus (*S_n_*_−1_) and the previous response (*R_n_*_−1_), merging all data from the current trial (**Fig. 1f,i**). In the absence of any history dependencies, *R_n_* would look identical for all conditional cases of *S_n_*_−1_ and *R_n_*_−1_. Otherwise, differences in *R_n_* indicate the presence of history-dependent influences, either taking the previous stimulus or the previous behavior into account.

In the grating motion experiments (**Fig. 1f**), larvae exhibited a higher-than-chance tendency to turn in the direction opposite the previous stimulus *S_n_*_−1_: after a rightward stimulus, in the next trial, they were more likely to turn to the left, and after a leftward stimulus, they were more likely to turn to the right, respectively. This tendency was present independently of the previous response. We will refer to this phenomenon as “stimulus repulsion” in the rest of the manuscript. At the same time, larvae exhibit a tendency to repeat the same behavior across trials. The probability of finding a left response was slightly higher after a previous left response, compared to when the previous response was right (**Fig. 1f**). We will call this phenomenon “response repetition”. We observed similar effects in the 100% coherence random-dot-motion experiments, with slightly weaker stimulus-history effects (**Fig. 1i**). These history-dependent effects were also observed across individual fish (**Fig. S1d,i**).

We then compared the strength of the sensory (**Fig. 1g,j**, left) and response (**Fig. 1g,j**, right). history effects across the two motion stimulation types. Again, for both grating and 100% coherence random-dot-motion stimuli, the previous stimulus direction had a negative effect, while the previous response had a positive influence on the current trial. The repetition effect of the previous response was weaker in the grating experiment compared to the 100% coherence random-dot-motion experiment (compare **Fig. 1g** and **Fig. 1j)**. We also repeated our analysis through an individual-animal-based quantification without bootstrapping, resulting in comparable effects (**Fig. S1c,d,h,i**). This analysis also showed that the previous stimulus had a repulsive effect on the current response, which was more prominent for the grating experiment than for the 100% random-dot-motion experiment (**Fig. S1e,j**).

This different behavior across motion types suggests that the strength of the stimulus influences which of the history phenomena dominates. Moving gratings have larger spatial structures without added stimulus uncertainty compared to the random-dot-motion cues. The short lifetime of the dots requires temporal integration (Bahl and Engert, 2020; Dragomir, Štih and Portugues, 2020), and animals may be more likely to rely on trial history in such experiments.

Taken together, these results show that larval zebrafish can be biased by previous motion trials that have occurred tens of seconds in the past. Previous stimuli have the tendency to induce repulsion, while previous behavior is generally repetitive. Such long-lasting traces of trial history can thus lead to incorrect responses in the current trial. As such, these effects may explain a certain fraction of trial-to-trial variability, as observed in many behavioral experiments under controlled laboratory conditions, without the need to postulate additional internal sources of noise.

### Previous evidence differentially influences stimulus and behavioral histories

Our observations that random-dot-motion stimuli, compared to gratings, induce a stronger behavioral repetition bias with weaker stimulus repulsion, suggest that history effects may be modulated by the degree of uncertainty in the stimulus. To investigate this idea further, we focused our experiments on the random-dot-motion stimulus, using four coherence levels, 0%, 33%, 67%, and 100%. In stimuli with lower coherence, the lower signal-to-noise ratio makes stimulus direction more challenging to extract. We designed experiments such that stimuli continuously moved for 30 seconds to the right or left from the perspective of the animal. To add a better perspective on the temporal scale of the sensory and behavioral histories, we used flexible inter-trial intervals between 0 and 60 s (**Fig. 2a**). Motion direction, coherence levels, and inter-trial intervals were randomized between trials.

**Figure 2.**
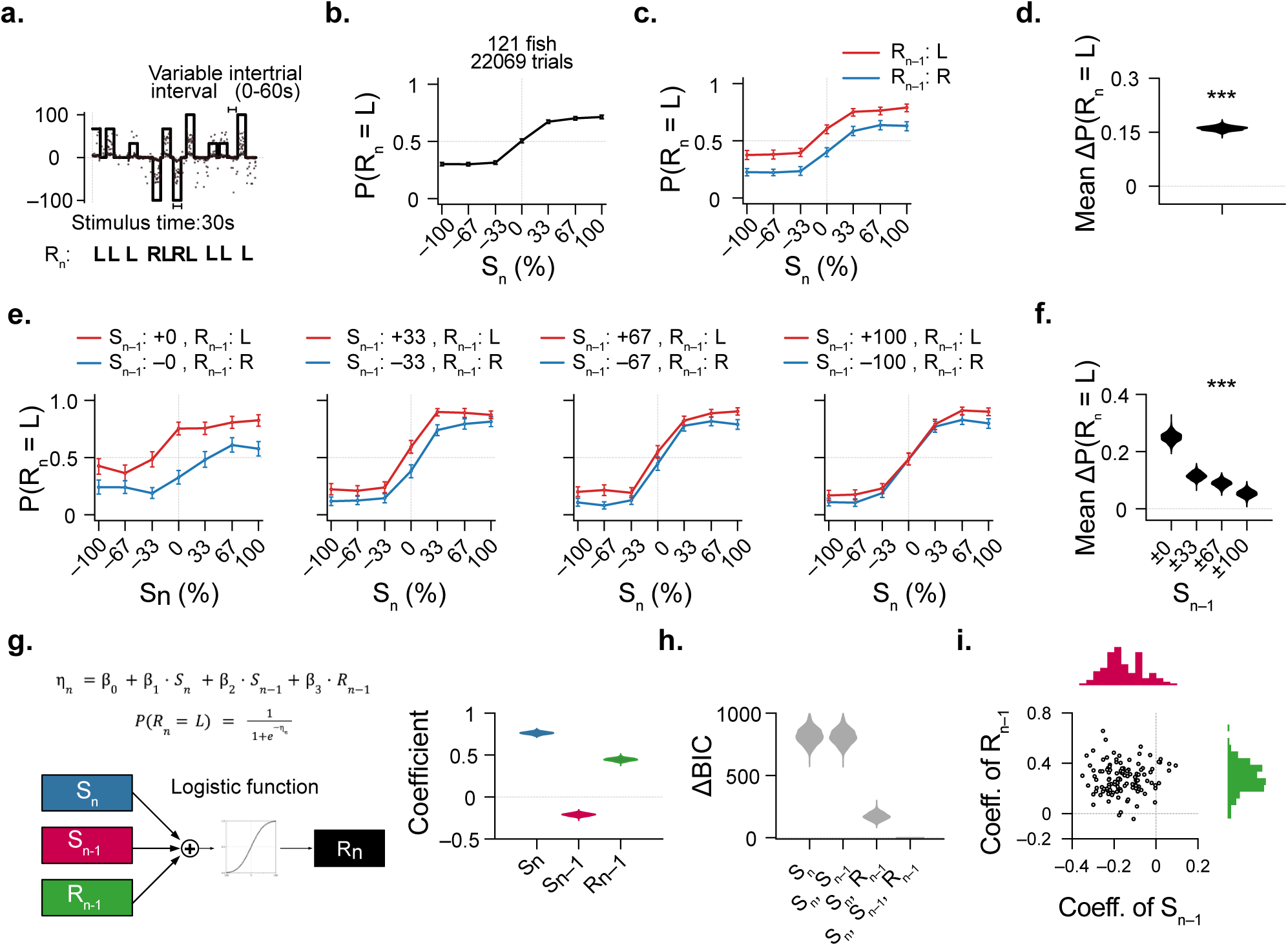
Stimulus and response histories have opposing and stimulus-strength-dependent effects on the current response. **a**, Schematic representation of the stimulus sequence of an example fish and bout turn angles (gray scatter). Responses are binarized into right and left, based on trial-based turn angle averages. **b**, Psychometric curve of proportion of trials with a left response. **c**, Psychometric curves conditioned on the response in the previous trial (*R_n_*_−1_). **d**, Bootstrapped distance of psychometric curves (*R_n_*_−1_ = *L* minus *R_n_*_−1_ = *R*) in (c). ***p < 0.001 when testing bootstrapped distances against zero (no difference between psychometric curves). **e**, Psychometric curves conditioned on the previous stimulus and the previous trial being "correct". We divided 0% trials into two groups (+0% and –0%) and assigned correctness labels randomly. **f**, Quantification of the average distance between the psychometric curves in (e). For each difference, ***p<0.001 (bootstrapping test against zero). ***p<0.001 for a monotonic decay as a function of *S_n_*_−1_ (permutation test against monotonic decay by chance, **Methods**). **g**, Generalized linear model with a logit activation function predicting the current response *R_n_* using the current stimulus (*S_n_*), the previous stimulus (*S_n_*_−1_), and the previous response (*R_n_*_−1_). **h**, Model complexity comparison using Bayesian information criteria, relative to the full three-parameter model in (g), ΔBIC. **i**, Hierarchical logistic regression model showing that relations are largely independent between the previous response and the previous stimulus coefficients for each fish (open circles). Spearman correlation between *S_n_*_−1_ and *R_n_*_−1_: *ρ* = 0.071, p = 0.446. N = 121 fish. See also related **Fig. S2**.

First, we sought to confirm results from previous studies on random-dot-motion evidence integration. As expected, we observed that response correctness increased (**Fig. 2b**) and that interbout intervals decreased (**Fig. S2a**, right) with stronger motion coherence levels. We also quantified performance for the first bout after stimulus onset and found that correctness increases with longer stimulus integration times (**Fig. S2b**). These results corroborate that larval zebrafish can integrate motion evidence during quiescence and that they can use such information to drive swimming. How does such behavior depend on the trial history? To test for this, we followed the analysis of our previous experiment (**Fig. 1**) by conditioning the psychometric curve on the previous response, regardless of whether the previous trial was left- or rightward (**Fig. 2c**). We found that when conditioned on the response in the previous trial being left (*R_n_*_−1_ = *L*), there was a higher tendency for leftward responses in the current trial. When conditioned on the previous trial being right (*R_n_*_−1_ = *R*), there was a higher tendency for rightward responses (**Fig. 2c**). To quantify these effects, we calculated the distance between the response-conditioned psychometric curves and found them to be around 15% (**Fig. 2d**). These results demonstrate that response repetition can significantly alter the optomotor response also for a stimulus with different coherence levels and variable inter-trial intervals.

Next, we asked how the strength of motion evidence biases responses. For this, we conditioned the psychometric curve on the previous trial (*S_n_*_−1_) to have a specified coherence level. To better disentangle motor and sensory history effects, we selected trials where the previous response direction matches motion direction, i.e., where the previous trial was labeled as correct. We found that the distance between previous-trial-conditioned psychometric curves decreases with increasing stimulus strength in the previous trial (**Fig. 2e,f**). Given our results, animals tend to repeat the same response across trials (**Fig. 2c**), the convergence of the psychometric curve for larger coherence levels indicates an increasingly negative effect of the stimulus strength, which counteracts the tendency to repeat behavior. Our analysis thus further supports the idea of two separate and opposing history-dependent forces to shape sensorimotor decision-making in the larval zebrafish.

To further quantify these effects, we employed a generalized linear model with a logit link function (same as performing a logistic regression; see **Methods**) to predict the current response from the current stimulus, the previous stimulus, and the previous response (**Fig. 2g**). The fitted coefficients of the predictors confirmed that the influence of the previous response and the previous stimulus are in opposing directions, with the positive effects of response repetition being slightly stronger than that negative effect of stimulus repulsion. We found a moderate positive correlation between the previous stimulus and the previous response (**Fig. S2c**), consistent with above-chance optomotor performance. High correlations between predictors can be problematic for generalized linear models because correlated predictors would compete to explain shared variance, making it difficult to attribute effects to one predictor versus another. We also modeled the dataset with fewer predictors and evaluated the model performance with the Bayesian information criterion (BIC). The BIC was at a minimum when we used the current stimulus, the previous stimulus, and the previous response as predictors, thus supporting the complexity of our proposed model (**Fig. 2h**).

So far, our analyses have been performed by collating trials across fish, followed by bootstrapping. To better understand the variability of history-dependent effects in the population, we employed a hierarchical logistic regression model to determine how the previous stimulus and the previous response predict for individuals (**Methods**). We found that more than 90% of fish exhibited the sensory- and response-dependent history effects (**Fig. 2i**), with the expected negative and positive coefficients, respectively. We could not find any significant correlation between these metrics across animals, suggesting that previous-trial stimulus repulsion and behavioral repetition use independent computational streams.

The variable inter-trial interval also allowed us to assess how long history-dependent effects may last. To this end, we grouped data into trials with short (0 to 25 s) and long intervals (35 to 60 s). For both groups, we found the positive response repetition effect to be indistinguishable, indicating that the motor memories can indeed last multiple tens of seconds without decay (**Fig. S2d**). Similarly, we analyzed the probability of repeating responses, dependent on stimulus strength (**Fig. S2e**). As we found earlier (**Fig. 2e,f**), response repetition was strongest for 0% coherence and decayed with higher coherence levels. Notably, these effects were independent of the inter-trial interval, suggesting that repulsive stimulus memories can also last multiple tens of seconds without decay.

Together, our experiments with variable inter-trial intervals and different random-dot-motion stimulus coherence levels provide further evidence that larval zebrafish sensorimotor decision-making is history-biased. The previous stimulus has a motion-strength-dependent negative effect, while the previous response induces a repetition bias.

### Increasing response repetition correlates with decreasing performance over the experiment

Our behavioral analyses and statistical modeling established that the previous response *R_n_*_−1_ and the previous stimulus *S_n_*_−1_ have opposing influences to shape the optomotor response in the current trial *n*. We next sought to examine the history-dependencies over multiple trials and over the entire experiment of three to four hours. To this end, we computed the autocorrelation of the response across trials, independently of stimulus direction. At a lag = 1 trial, we obtained an autocorrelation value of 7%, corroborating a strong trial-to-trial behavioral repetition effect (**Fig. 3a, Fig. S3a)**. In history-dependency studies in humans and rodents, values are in the order of 5 to 10% (Fischer and Whitney, 2014; Hachen *et al*., 2021, 2026). The autocorrelation function then decayed slowly with an exponential decay constant of 15 trials (**Fig. 3a**, **Fig. S3b).**

**Figure 3.**
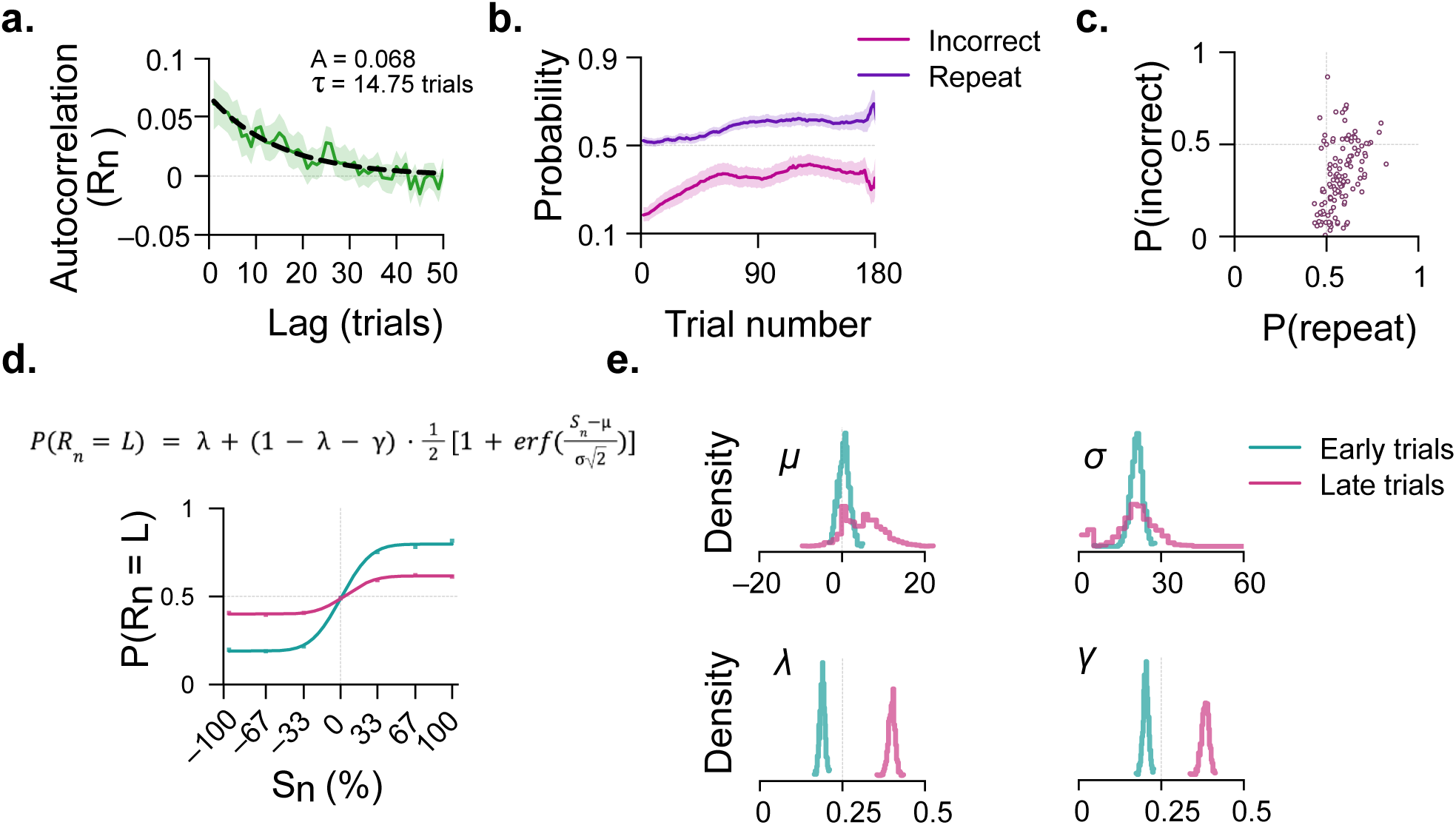
Response repetition increases over the course of the experiment. **a**, Autocorrelation function of binarized trial responses. **b**, Probability of response repetition and response incorrectness as a function of experimental trial number. Median Spearman correlation between P(repeat) and trial number was 0.061, p < 0.001 (Wilcoxon signed-rank test against zero). Median Spearman correlation between P(incorrect) and trial number was 0.104, p < 0.001 (Wilcoxon signed-rank test against a Spearman correlation value of 0). **c**, Correlation between response repetition and trial incorrectness rate. Each circle represents the means of one fish. Spearman correlation was 0.503, p < 0.001. **d**, Psychometric curves, grouped by early (within the first 70 minutes of the experiment) and late (after 120 minutes into the experiment) trials. Circles represent bootstrapped means. Solid lines represent fits using the cumulative Gaussian error function (equation on top and **Methods**). **e**, Normalized distributions of the fitted model parameters of inflection point (*Δμ* = 4.446 %, p = 0.34), sensitivity (*Δσ* = −0.668 %, p = 0.99), and the asymptotes (lower asymptote, *Δλ* = 0.21, p < 0.001; upper asymptote, *Δγ* = 0.18, p < 0.001). Bootstrapped test of the differences (**Methods**). Data from N = 121 fish (same as in Fig. 2). Solid lines in (a,b) indicate means of bootstrapped data. Transparent shading in (a,b) indicates 95% confidence intervals of bootstrapped probabilities. Also see related **Fig. S3**.

We next sought to explore how stable these effects are over the time course of the experiment. To answer this, we computed the probability of response repetition, *P*(*repeat*), over trials (**Fig. 3b, Methods**). We found that *P*(*repeat*) increases over the course of the experiment. This increasing tendency to repeat responses in uncorrelated stimulus sequences should lead to decreasing optomotor performance as the trial number increases. Consistent with this expectation, we found an increase of incorrect trials, *P*(*incorrect*), over time **(Fig. 3b)**. Correlating *P*(*repeat*) and *P*(*incorrect*) across fish displayed a strong positive correlation between these metrics (**Fig. 3c**), suggesting that response repetition plays a major role in shaping optomotor performance.

To ensure that this reduced optomotor response is not a symptom of decreasing activity over time, we also computed the interbout interval and bout distance over the experiment. If the fish were to become less active, interbout intervals should rise, and the bout distance should decrease. Contrary to this, we found that the interbout interval decreased slightly and bout distance increased over time **(Fig. S3c,d)**, suggesting that the observed decrease in optomotor performance originates from altered sensorimotor processing and not activity levels.

To identify the reasons behind the observed increase in incorrect trials, we modeled behavior with a classical psychometric curve as a cumulative Gaussian error function with four parameters (Wichmann and Hill, 2001) (**Fig. 3d**, **Methods**). In this framework, the increase in incorrect trials over the experiment can be explained in two hypothetical ways that make different predictions about the shape of the psychometric curve and its parameterization: 1) Larvae could become worse at discriminating the stimulus at the sensory level. A decrease in stimulus discrimination ability would flatten the curve and decrease the sensitivity (*σ*) parameter, since small changes in coherence would be harder to distinguish. 2) Larvae could increasingly respond independently of the current stimulus, also known as lapse rate in psychophysics. Stimulus-independent responses, on the other hand, would change performance for even the strongest stimulus such that the lower and upper asymptotes of the curve change toward chance levels. This effect is modeled by adjusting the lower (*λ*) and upper (*γ*) lapse rate. The *μ* parameter describes the horizontal shift of the curve and relates to the inflection point or the point of subjective equality. Such an effect would indicate an altered swimming bias, which is unlikely to occur in our symmetric experimental design.

To address which of these processes significantly influence the decrease in performance over time, we estimate psychometric parameters in data from early and late trials (within the first 70 minutes and after 120 min of the experiment, respectively) (**Fig. 3d**). We found that late trials have substantially increased lower and upper lapse rates compared to early trials, whereas the sensitivity and the horizontal shift parameters remained unchanged (**Fig. 3e**), as proposed in hypothesis 2, mentioned above.

Thus, our results reveal an increased response repetition over experimental time that largely influences the rate of incorrect trials. By extracting psychometric parameters from animals in early and late trials, we find that lapse rates increase over time while we do not observe significant changes in sensitivity. Together, our findings suggest that repeating responses may be a primary cause for the observed bypassing of current sensory information during sensorimotor decision-making.

### A deterministic multi-scale integrator model captures history-dependent behavioral variability

In the previous sections, we showed that larval zebrafish responses to motion stimuli are biased by stimuli and behavior in the recent past. To understand the mechanism by which history biases propagate across trials, we modeled the sensorimotor pathway as three parallel leaky integrators, each evolving continuously in time with its own decay time constant (**Fig. 4a**). Two integrators operate on the stimulus, while another one integrates motor responses over time. To prevent the self-reinforcing motor integrator from diverging, we limited motor integration to stimulation periods so it has time to decay in between. Based on our finding that a simple general linear model with three linear predictors can describe trial-to-trial history effects (**Fig. 2g**), we designed the model output to be a weighted sum of the integrated values. The model then generates turn angles as a continuous function over time. We tuned parameters based on previous literature (Bahl and Engert, 2020; Dragomir, Štih and Portugues, 2020; Slangewal *et al*., 2026) and using a systematic heuristic approach (**Methods**).

**Figure 4.**
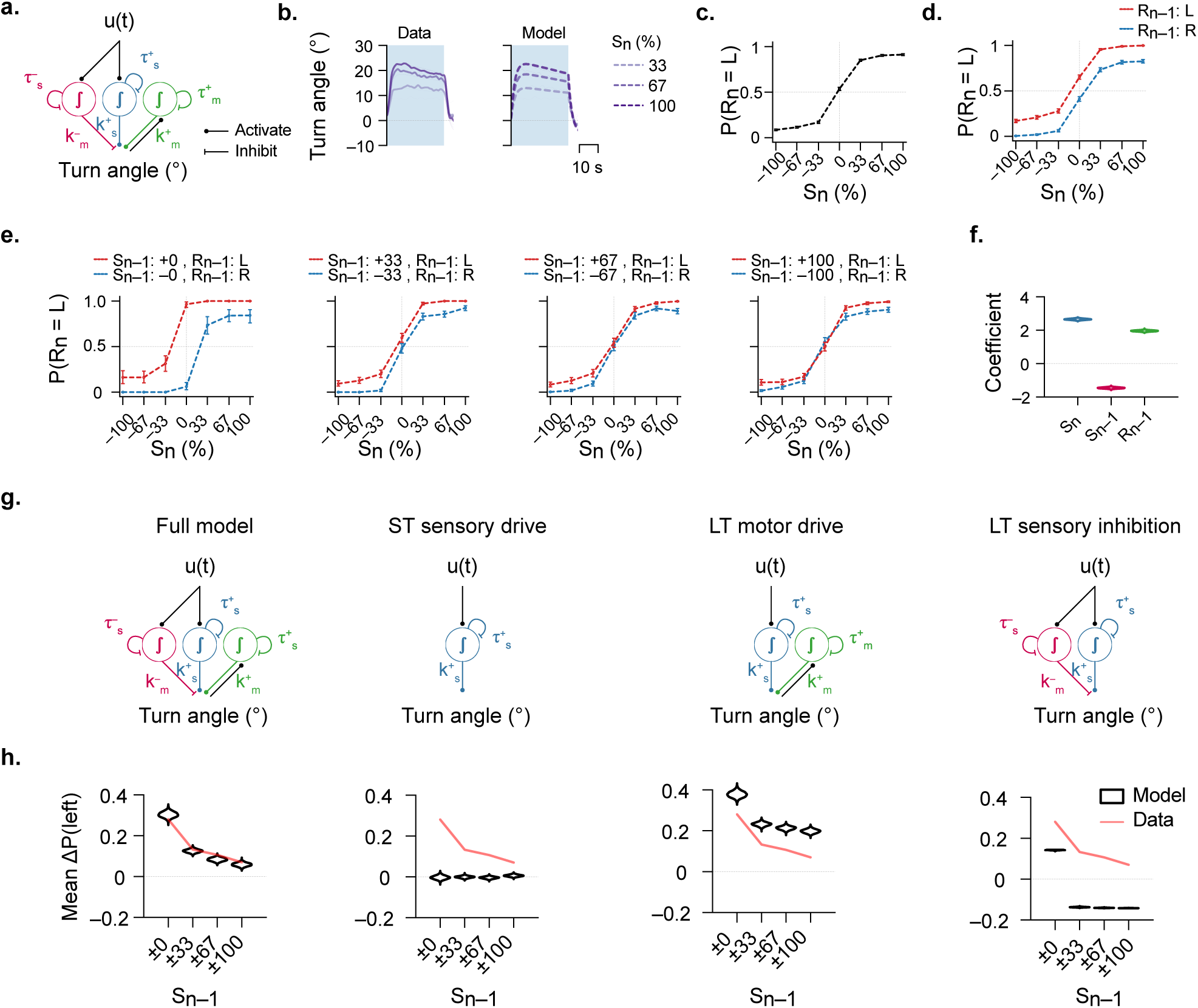
A deterministic multi-scale integrator model captures history-dependent behavioral variability. **a,** Schematic of the multi-scale integrator model with three leaky integrators operating on sensory and motor streams across different temporal scales (*τ*). Integrator outputs are weighted and summed to generate turn angles (**Methods**). **b**, Within-trial response dynamics of data (left) and model (right). **c,** Psychometric curve showing performance of the model operating on the same trial sequences as used in our experiments (for example, Fig. 2a). Dashed lines and dots represent mean bootstrapped simulations, as in Fig. 2b. **d,** Psychometric curve of the model, conditioned on the previous response. **e,** Psychometric curves, conditioned on the previous trial having a correct response and specified coherence levels, as in Fig. 2e. **f,** Generalized linear model coefficients fitted to model output, as in Fig. 2g. **g,** Representation of tested model variants. LT = long-term; ST = short-term. **h,** Quantification of distance between psychometric curves for each model variant. For the full model, this is a quantification of panel (e). Bootstrapping done as in Fig. 2f. The red line indicates the means of the experimental data shown in Fig. 2f. Model simulations on N = 121 different stimulus streams identical to the ones used in the experiments in Figs. 2 and 3. Also see related **Fig. S4**.

Notably, the model has no noise in sensorimotor transformation and is fully deterministic in its implementation. This configuration allows us to probe to what extent sensorimotor trial history can shape behavior across the experiment. To probe the model with experimentally realistic conditions, we chose the same trial sequences (including stimulus time and inter-trial intervals) as we had used for real larvae.

We first wanted to know if our model can capture behavior on shorter time scales. To this end, we followed previous quantification approaches (Slangewal *et al*., 2026) that analyze turn angle within a trial. The model captures onset dynamics and an apparent decrease over the stimulus period (**Fig. 4b**). We attribute these features to the positive and negative sensory integrators (**Fig. 4a**, blue and pink, respectively) having different time constants. Using our experimental data with gratings and 100% random-dot-motion (**Fig. 1**), we further analyzed dynamics within the inter-trial period. Our model predicted some degree of swimming in the opposite direction of the previous stimulus (**Fig. S4a**). We observed dynamics to be qualitatively similar in our data for both motion stimulus types. We therefore suggest that the slow negative sensory integrator can carry over into the next trial, which may be the source of the observed repulsive bias.

Given that our model can carry over biases from the previous trial, we next sought to probe if it can capture the key features of the psychometric curves we experimentally measured before (**Fig. 2**). Even though our model behaves deterministically, we observed a large (∼17.2 %) fraction of trials to be incorrect, also for strong stimuli (compare **Fig. 4c** and **Fig. 2b**). When we condition our model results on the previous response (*R_n_*_−1_; **Fig. 4d** and **Fig. S4b**) or on both previous stimulus and response (*S_n_*_−1_ and *R_n_*_−1_; **Fig. 4e**), we recover results qualitatively matching our experimental results (**Fig. 2c–e**).

To find the overall contribution of the integrators to the behavior, we subjected the model output to a generalized linear model, as before (**Fig. 2g**). Although correlations between predictors were higher than in the data (compare **Fig. S4c** and **Fig. S2c)**, values were still small enough to allow for coefficient estimation in a generalized linear model. Moreover, model comparison using BIC shows that including all three predictors provides the best explanation of the model output (compare **Fig. S4d** and **Fig. 2h**). We recover comparable contributions of the individual predictors (*S_n_*, *S_n_*_−1_, and *R_n_*_−1_; compare **Fig. 4f**, **Fig. 2g**, and **Fig. S4e)**.

Next, we asked if the proposed complexity of our model is indeed required to capture the observed history-dependencies. We addressed this question by generating simpler model variants by taking our full model and setting the gains of the repulsive integrator (pink) and/or the motor integrator (green) were set to zero (**Fig. 4g**). We name model variants based on the components they possess: full, short-term (ST) sensory drive, long-term (LT) sensory inhibition and LT motor drive (**Fig. 4g**). We then analyzed and compared the outputs of these models in the same manner as before (**Fig. 2e,f**).

The ST sensory drive model and the LT motor drive model fail to reproduce the slow decay of turn angle within the stimulus period **(Fig. S4f,g)**. We attribute this observation to the missing negatively weighted slow integrator (pink). Quantifying the distance between stimulus- and response-conditioned psychometric curves (**Fig. 4d,e**), as in **Fig. 2e,f**, we find that all model variants, but the full model, fail to accurately reproduce the data. As expected, the ST sensory drive model behaves effectively memoryless (**Fig. 4g**). The LT motor drive could only qualitatively capture effects. The LT sensory inhibition model displayed an apparent response repetition for *S_n_*_−1_ = 0% coherence, even though it does not have a positive motor integrator. We explain this observation as follows: *R_n_*_−1_ depend on repulsive memory arising from the stimulus in trial *n* − 2. Since *S_n_*_−1_ = 0 % does not alter responses, the resulting response differences carry to the current trial. In cases where *S_n_*_−1_ > 0%, the repulsive effect of trial *n* − 1 dominates history effects.

In summary, our multi-scale integrator model can explain trial-to-trial history effects, while being completely deterministic without any stochastic processes. To capture short (within trials), intermediate (during inter-trials), as well as trial-to-trial behavioral dynamics, we need a combination of a negative and a positive stimulus integrator together with a positive motor integrator that operate at different temporal scales. As such, our results show that the sequence and history of stimuli and responses can considerably impact behavioral variations and may well explain a fraction of incorrect trials in our experiments.

### The multi-scale integrator model captures experiment-level dynamics

Finally, we asked whether the model could also account for the longer-timescale phenomena observed in the data. To test this, we compared model output to three key experimental results: 1) The response repetition increases over time (**Fig. 3b**). 2) The probability of incorrect responses increases over time (**Fig. 3b**). 3) The early and late trials display different parameterization of the psychometric function (**Fig. 3d,e**).

All three effects are reproduced in the simulations using our full model (**Fig. 4a**) with unchanged time constant and gain values (**Fig. 5a,b)**. The ability of our model to explain the increase of response repetition and incorrect trials over experimental time comes from the self-enhancing motor integrator. This integrator operates on the timescale of around 15 trials (**Fig. 3a**), and can thus lead to a slow accumulation of motor bias.

**Figure 5.**
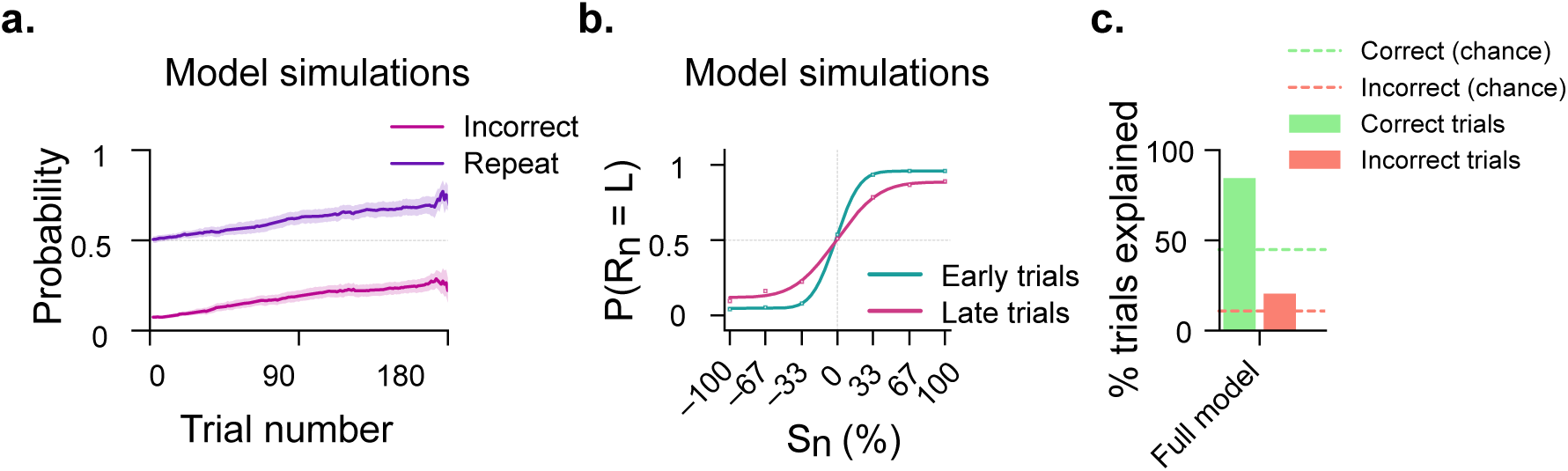
The multi-scale integrator model captures experiment-level effects. **a,** Model simulations showing probabilities of repeating a response from trial to trial and of incorrect responses over trials in the experiment. Median Spearman correlation of repetition over time = 0.1; p < 0.001, Wilcoxon signed-rank test against 0; Median Spearman correlation of incorrectness over time = 0.143; p < 0.001, Wilcoxon signed-rank test against zero.**b,** Early (less than simulated 70 min) and late (after simulated 120 min) psychometric curves from model simulations. **c,** Percentage of correct and incorrect trials above chance explained by the model. Chance levels of labelling a trial “correct” or “incorrect” based on the rate of correct trials in the data. Model simulations on N = 121 different stimulus streams identical to the ones used in the experiments in Figs. 2 and 3. Also see related **Fig. S5**.

When analyzing the parameterization of the psychometric curves (**Fig. S5a**), we observed an increase in lower (*λ*) and upper (*γ*) lapse rates, and a similar inflection point (*μ*), as in our data (**Fig. 3e**). In our model simulations, the sensitivity parameter *σ* was larger in late than in early trials, an effect we did not observe in our experiments. We speculate that this discrepancy is due to additional behavioral variability in real fish (Krishnan *et al*., 2025) beyond stimulus and motor history effects.

We also performed analyses in the reduced model variants (**Fig. 4g**). As expected, models without a motor integrator failed to capture experiment-level dynamics (**Fig. S5b–d**, ST sensory drive and LT sensory inhibition models, left and right, respectively). A missing negative stimulus integrator, however, did not affect these long time scales (**Fig. S5b–d**, LT motor drive model, middle). Only the LT motor drive model succeeded in qualitatively capturing the shape of the psychometric curve, including the asymptotes for strong motion stimuli (**Fig. S5c,d**, middle).

As our models have all been applied to the same stimulus sequences as used for the experiment, we next quantified to what extent model predictions and experimental data match (**Fig. 5c**). This analysis allows us to probe how much of the observed behavioral variability can be explained by history-dependent sensorimotor processes. We first computed the overall experimentally observed rate of correct (∼67 %) and incorrect trials (∼33 %). Randomly drawing from such rates to predict trial outcomes correctly would thus give chances of ∼45 % to predict correct trials as correct and ∼11% to predict incorrect trials as incorrect (dashed lines in **Fig. 5c**). Our full model predicted around 85% of the correct trials to be correct and around 20% of the incorrect trials to be incorrect, a major improvement over chance levels. Our alternative model lacking repulsive stimulus memory (LT motor drive) could capture these effects as well. Model variants without a motor integrator (ST sensory drive and LT sensory inhibition models) predicted an even larger fraction of correct trials, which is expected as they deterministically take into account mostly the momentary stimulus. Yet, these models failed to improve predictions of incorrect trials beyond chance.

Our modeling and simulation results demonstrate that the multiple integrator model robustly captures correct responses while also accounting for error-generating processes such as hysteresis. Critically, while each reduced model variant could reproduce only a subset of these phenomena, only the full model, with a single, unchanged parameter set, jointly captured effects spanning stimulus-driven, trial-to-trial, and experiment-level timescales, highlighting motor memories as a major source of behavioral variability.

## DISCUSSION

In this study, we investigated the impact of trial history on sensorimotor processing in the larval zebrafish. By applying classical psychophysical methods (Green and Swets, 1966; Wichmann and Hill, 2001; Carandini and Churchland, 2013) to an optomotor paradigm (Orger *et al*., 2000, 2008; Portugues and Engert, 2011; Bahl and Engert, 2020; Dragomir, Štih and Portugues, 2020; Garza, Hady and Bahl, 2026), we found that trial history considerably modulates responses to incoming sensory information. Our findings show that sensory history is repulsive and response history is attractive. Responses are pushed away from the direction of the previous stimulus, while they are pulled towards the previous response. For example, a leftward stimulus increases the probability of rightward responses in the next trial. At the same time, a leftward response will increase the probability of leftward responses. Such a pattern of opposing influences of sensory and decision history has been reported in other vertebrates, including humans (Lages and Treisman, 2010; Fritsche, Mostert and Lange, 2017; Pascucci *et al*., 2019; Bosch *et al*., 2020).

One possible cause for repulsion may be neuronal adaptation in early sensory processing stages, reducing sensitivity (Magnussen and Johnsen, 1986; Pascucci *et al*., 2019). For example, on prolonged exposure to a directional moving stimulus, humans report perceiving movement in the opposite direction. This phenomenon is known as the motion aftereffect or the waterfall illusion (Gibson, 1937; Biber and Ilg, 2011). In addition, humans display eye movements in the opposite direction to previous stimulation (Anstis, Verstraten and Mather, 1998; Mather *et al*., 2008). In the larval zebrafish, the motion aftereffect has been shown for eye movements and tail flicks (Pérez-Schuster *et al*., 2016; Wu *et al*., 2020). While the motion aftereffect may partially contribute to the repulsion in our dataset, our results suggest that it is unlikely to be the main driver: First, previous work on the motion aftereffect in the larval zebrafish showed that the stimulation duration must be at least multiple minutes (an order of magnitude longer than the stimulus duration used in the present work) to evoke an observable aftereffect (Pérez-Schuster *et al*., 2016; Wu *et al*., 2020). Second, neuronal adaptation at the early sensory level has rapid recovery times (Greenlee *et al*., 1991), with aftereffects lasting a fraction of the stimulation duration, whereas the repulsion we observed scales beyond the timescale of stimulus presentation. An additional argument against the role of early sensory adaptation in the observed repulsive behaviors is our measured psychometric curves, which we conditioned on the previous stimulus. Our analyses show a change in bias rather than in sensitivity, suggesting that sensory repulsion occurs at a later processing stage than sensory encoding (Wichmann and Hill, 2001; Carandini and Churchland, 2013). To this end, previous work in mammals explained repulsive effects as the trial-by-trial modulation of evidence accumulation and decision-making processes (Treisman and Williams, 1984; Hachen *et al*., 2021; Schönsberg *et al*., 2025).

In rodents and humans, response repetition can be explained by reinforcement mechanisms induced by rewarding correct decisions (Lak *et al*., 2020). Repetition effects are particularly strong after trials with ambiguous sensory information. In these animals, decision-making experiments involve training over several trials or sessions. Our findings show that strong repetition can also be found in behavioral paradigms without explicit reward and training, suggesting the possibility that response repetition exists as an innate and learning-independent mechanism.

In studies of trial history, an important factor to consider is that the stimulus and response are correlated, making it challenging to identify the source of hysteresis. Serial dependence in humans has been attributed to sensory history (Fischer and Whitney, 2014) as well as response history (Fritsche, Mostert and Lange, 2017; Bosch *et al*., 2020). Such discrepancies could arise from task design or choice of analysis methods. In larval zebrafish, it has been reported that sequentially repeated stimuli in both optomotor and obstacle avoidance tasks lead to enhanced performance (Tanaka and Portugues, 2025; Zhao *et al*., 2026). Both studies attribute this effect to stimulus history rather than behavioral history. Our analysis based on conditioning psychometric curves on sensory or response history allows us to disentangle effects. We argue that sensory history is actually repulsive, but that this negative bias is overcome by behavioral history promoting response repetition in consecutive trials. The differently designed behavioral paradigms may also contribute to the discrepancy between our and previous interpretations. While we probe responses of freely swimming zebrafish to sideward moving optic flow, the other works used head-restrained animals responding to rotational and front-to-back visual cues.

We show that history-dependent biases are carried over from trial to trial and accumulate over hours. Based on these observations, we built a deterministic model that captures behavioral results across temporal scales and that can explain a considerable fraction of correct and incorrect trials. We also observed that the optomotor performance decreases over experimental time. The psychophysical signature of this reduction was an increase in lapse rates, a phenomenon typically attributed to attentional fluctuations (Wichmann and Hill, 2001; Ashwood *et al*., 2022). Consistent with previous work in rodents (Pisupati *et al*., 2021; Gupta *et al*., 2024), we also observed that trial-history biases can lead to apparent lapses in our dataset. In larval zebrafish, it has recently been suggested that disengaged states may be a source of optomotor variability (Krishnan *et al*., 2025). Our results complement these findings: Repetition-driven hysteresis can produce behavioral variability without invoking attentional disengagement or stochasticity in sensorimotor processing.

In mammals, representations of trial history in the brain have been found in various regions of the cortex in both rodents (Akrami *et al*., 2018; Findling *et al*., 2025; Hachen *et al*., 2026) and primates (Rao, DeAngelis and Snyder, 2012). Neurons with slow dynamics matching our observed behavioral timescales have been found in the torus longitudinalis of the larval zebrafish midbrain (Bahl and Engert, 2020). This region could be a potential neural substrate for mediating history effects in sensorimotor processing. In addition, the activity of the anterior rhomboencephalic turning region (ARTR) has been hypothesised to control streaks of bouts in the same direction and may underlie behavioral repetition across trials (Dunn *et al*., 2016). The dorsal thalamus may also be an important structure in this context, as it has recently been shown to hold a persistent representation of previous sensory stimuli (Zhao *et al*., 2026). Our model suggests that in the absence of optic flow, the motor integrator remains inactive, a property required to prevent swimming from being locked indefinitely in a certain direction. Potential candidate regions in the larval zebrafish brain, are the cerebellum, torus longitudinalis, and inferior olive being (Ali *et al*., 2023; Narayanan, Varma and Thirumalai, 2024; Tanaka and Portugues, 2025).

The sensory motor history features we have identified in larval zebrafish optomotor behavior across broad temporal scales may relate to the dynamic features of the environment in which animals navigate (Zhao *et al*., 2026). Thus, our findings hint at potential links between such mechanisms and predictive sensory processing (Pouget *et al*., 2013; Narayanan, Varma and Thirumalai, 2024; Findling *et al*., 2025).

In conclusion, we report that opposing and long-lasting influences of sensory and response history in the larval zebrafish are qualitatively similar to what has been reported in animals with cortical structures such as rodents and humans (Fritsche, Mostert and Lange, 2017; Bosch *et al*., 2020; Hachen *et al*., 2021). Our deterministic model can explain a considerable fraction of trial-to-trial variability, showing that history effects have to be taken into account when interpreting behavioral data. Using larval zebrafish, an animal with molecular and neural access, for such analyses holds promise to access hypothetically conserved neural circuit mechanisms underlying persistent sensorimotor computations.

## METHODS

### Zebrafish rearing

Adult wild-type zebrafish (*Danio rerio)* of AB strain and KN strain (in grating experiments, **Fig. 1**) were used to generate larvae for experiments. Eggs were collected from crosses in E3 water with methylene blue. After one day, the water was replaced with fresh E3 water without methylene blue. Eggs and larvae were raised in Petri dishes with a diameter of 14.5 cm in an incubator set to a 14h:10h light-dark cycle. For all fish used in **Fig. 1**, the incubator temperature was stably maintained at 28 °C. For fish used in **Figs. 2** and **3**, the incubator temperature was stably maintained at 28 °C during the daytime. All experiments were performed on 5- to 7-day-old larvae. All experiments done on larvae older than 5 dpf were approved by Regierungspräsidium Freiburg, Germany, under permit number G21/153 (“Soziales Lernverhalten im larvalen Zebrafisch”).

### Behavioral experiments

Zebrafish larvae were placed in custom-made 6-cm-radius circular arenas with black acrylic rims and translucent floors. Each arena was placed under a camera (Basler acA2040-90um-NIR) equipped with a zoom lens (Navitar ZOOM 7000) and an infrared filter (850 nm, longpass) and over infrared LED lights (940 nm, SOLAROX). Projectors (AAXA P300 Pico Projector) were used to present the stimulus on the arena floors from below. Floors were coated with diffusive paper inside the water. The cameras were operated at 90 Hz and the projectors at 60 Hz.

Zebrafish larvae were tracked in real time using custom-written Python 3.12 software (Capelle *et al*., 2026). The centroid of the largest tracked blob after background subtraction was used to determine the position of the larvae. The body orientation was computed by using body shape and real-time posture analysis, followed by principal component analysis. Accumulated orientation was calculated as the cumulative orientation change of the animal. To detect bouts, a 50 ms rolling variance of the accumulated orientation was calculated. A bout was defined to start when this value crossed 1 deg^2^ for at least 20 ms. The bout was defined to have ended when the rolling variance fell below 0.5 deg^2^ for at least 50 ms. For each bout, start and end positions, duration, turn angle, distance travelled, and interbout interval were extracted.

Stimuli were delivered using the same custom-written software framework. Larval zebrafish were presented with continuous whole-field leftward or rightward visual motion. The direction of motion was locked to the body orientation of the freely swimming fish in real-time to maintain an identical signal for the duration of the stimulus. Motion was presented for 30 seconds with an inter-trial interval of 40 seconds.

In the grating experiment (**Fig. 1e–g**), we used moving alternating white and black stripes whose luminance varied sinusoidally. The stimulus moved at 1.8 cm/s and had a spatial wavelength of 1.2 cm. A grey homogeneous background was presented in the inter-trial period. We measured arena luminance values using an LX1330B light meter (Hongkong 854 Thousandshores Ltd., Central Hong Kong) held around 25 cm above the arena: Grating stimulus was about 85 lux, the gray periods was about 75 lux, and all random-dot-motion stimuli was about 20 lux.

In the 100% dot motion experiments (**Fig. 1h–j**), 1200 white dots were presented on a black background. All dots moved with a speed of 3.6 cm/s but were redrawn at a random position after an average lifetime of 200 ms. 0% coherence was presented in the inter-trial period, during which dots were only randomly redrawn. The variable inter-trial random-dot-motion experiments (**Fig. 2**) used the same parameters, except with varying inter-trial intervals and coherence levels (fraction of dots moving). We used seven coherence levels: ±0%, ±33%, ±67%, and ±100%. Positive values indicate motion to the left, and negative values indicate motion to the right. We showed 0% twice to generate the same amount of data for our response-history-based analyses (**Fig. 2e**, left). The order of stimuli was randomized, and all stimuli were equally likely to be presented.

### Behavioral analysis

Bouts were considered for analysis only if they fulfilled the following criteria: 1) interbout intervals smaller than 10 s, 2) contour area of the detected fish larger than 50 and smaller than 1000 pixels, 3) absolute turn angle smaller than 150 degrees, and 4) radial position smaller than 5.75 cm. This filtering was done to ensure that tracking air bubbles, small scratches on the arena, interactions with the wall, or swapping of head and tail did not impair our analysis. We only included trials in which all of these criteria were fulfilled for at least 80% of bouts. We considered only fish where at least 80% of the trials were valid. With this filtering strategy, we obtained 160 fish and 15981 trials in experiments with sine gratings (**Fig. 1e–g**), 76 fish and 7416 trials in experiments with 100% dot motion (**Fig. 1h–j**), and 121 fish in experiments with mixed coherence dot motion (**Figs. 2** and **3**).

Trial-averaged turn angles (*Δα*) were computed by first averaging the turn angle of all bouts in a trial (during motion presentation). This value was then binarized: left response (*R_n_* = 1), when *Δα* > 0°) or right response (*R_n_* = 0, when *Δα* < 0°).

We used a moving window average to compute *P*(*repeat*) and *P*(*incorrect*) over experiment time **(Fig. 3b)**, with a forward-looking window of size 30 trials and a step size of 1 trial.

Within-trial dynamics were quantified using a moving window average on turn angle (**Fig. 4b**). Specifically, the averages were computed with a backward-looking window size of 1 s and a step size of 0.1 s. For better visualization, we smoothed the mean using scipy’s *gaussian_filter1d* function with sigma = 5 bins (**Fig. 4b**, left).

### Statistical analysis

Unless stated otherwise, statistical significance was computed by bootstrap resampling of trials pooled across fish. We picked the same number of trials as in the complete dataset with replacement 1000 times. All resampling was done by stratified bootstrapping, that is, the dataset was first divided based on the conditioning, and then each subset was resampled. We show means with 95% confidence intervals of the bootstrap distributions.

In **Fig. 1g,j**, statistical significance in *S_n_*_−1_ and *R_n_*_−1_ effect between experiments was tested by first building a difference-of-bootstrap-distributions (1000 iterations per stimulus type and subtracted element-wise) and then computing a two-sided bootstrap p-value against a null-distribution generated by shuffling the data.

In **Fig. 2f**, statistical significance of monotonic decrease was assessed using a permutation test. For each of 1000 bootstrap iterations, we obtained one value per condition (*S_n_*_−1_ = ±0, ±33, ±67, ±100) and counted the number of pairwise inversions relative to a strictly monotonic decreasing sequence; these counts were averaged across bootstrap draws to yield the observed test statistic. A null distribution was generated by, for each of 10000 permutations, randomly assigning the correspondence between values and conditions within each bootstrap draw and recomputing the mean inversion count. The p-value was defined as the fraction of permutations yielding a mean inversion count less than or equal to the observed value.

In **Figs. 2i, 3b,c**, **S3c,d**, **5a**, and **S5b**, Spearman correlations were computed using the *spearmanr* function in the *scipy.stats* library in Python. The Wilcoxon signed-rank test was performed using the *wilcoxon* function from the same library.

In **Figs. 3d** and **5b**, we classified trials as early if they occurred within the first 70 minutes of the experiment and as late if they occurred after 120 minutes into the experiment. Psychometric curves in **Fig. 3d** were fitted to the cumulative Gaussian error function using the *curve_fit* function from *scipy.optimize*:

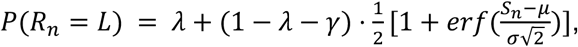

where *μ* describes the horizontal shift, *σ* relates to the sensitivity, and *λ* and *γ* represent the lower and upper lapse rates, respectively.

### Generalized linear model

From the binarized response dataset, we aimed to predict the current response (*R_n_*) using three predictors (**Fig. 2g**): the current stimulus (*S_n_*), the previous stimulus (*S_n_*_−1_), and the previous response (*R_n_*_−1_). We used a generalized linear model with a logistic link function using *sklearn’s* LogisticRegression framework:

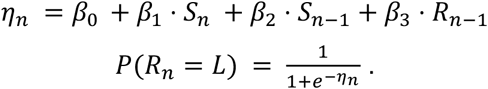

We used “l2” penalty, “lbfgs” solver, and C=1 for regularization. These parameters were chosen to ensure numerical stability across bootstrap iterations. We tested four models (**Fig. 2h**): 1) only *S_n_*, 2) *S_n_* and *S_n_*_−1_, 3) *S_n_* and *R_n_*_−1_, 4) *S_n_*, *S_n_*_−1_ and *R_n_*_−1_. To quantify variability and robustness of the parameters, we used a bootstrap approach. In each of the 1000 iterations, we sampled trials with replacement and fit the model to the resulting bootstrap sample. We evaluated the model performance using the Bayesian Information Criterion (BIC).

### Hierarchical logistic regression model

For estimating variability of history dependencies across fish, we implemented a hierarchical logistic regression model using the PyMC (Abril-Pla *et al*., 2023) and ArviZ (Martin *et al*., 2026) libraries (**Fig. 2i**). As in the generalized linear model, the hierarchical model predicts the probability of a leftward response *R_n_* based on three predictors: current stimulus (*S_n_*), previous stimulus (*S_n_*_−1_), and previous response (*R_n_*_−1_).

At the population level, we used weakly informative priors on all regression coefficients (*μ_Sn_*, *μ_Sn_*_(*_, *μ_Rn_*_(*_) and intercept (*μ_α_*) by drawing from a normal distribution *N*(0, 1.5). The corresponding standard deviations (*σ_Sn_*, *σ_Sn−1_*, *σ_Rn−1_*, *σ_α_*) relate to inter-individual variability and were drawn from the positive side of a normal distribution *N*(0, 1.0). Fish were treated as random effects, such that each fish *i* had its own set of coefficients drawn from these population-level distributions:

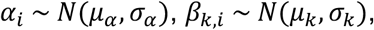

where *k* ∈ {*S_n_*, *S_n_*_−1_, *R_n_*_−1_}. This hierarchical structure enables partial pooling, allowing fish-specific estimates to be regularized toward the population means, depending on the amount of data available per fish. The probability of a leftward response was modeled using a logistic link function with fish-specific coefficients, using the same formulation as for the generalized linear model (above).

### Multi-scale integrator model

We modeled the turn angle of the fish (*y*(*t*)) as a continuous-time system with three parallel integrators. The model processes the trials in the same sequence and temporal arrangement as experienced by the fish. The model stores average turn angles during the stimulus period in the same manner as we do for real fish, enabling identical analysis of model and experimental data.

The model implementation is as follows:

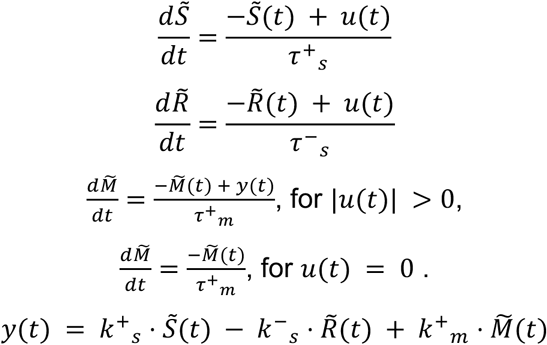

Here *u*(*t*) describes the stimulus input, with 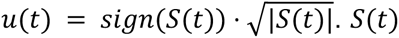 is the coherence level (scaled between –1 and 1) during the trial and inter-trial period. *S̃*, *R̃*, and *M̃* are the attractive and repulsive stimulus integrators, and the attractive motor integrator, respectively. *k*^+^*_s_*, *k*^−^*_s_*, *k*^+^*_m_* are the weights that combine the different integrators into model output *y*(*t*). We clip *M̃*(*t*) between –1 and 1 and *y*(*t*) between –50 and 50 to ensure numerical stability. Simulations were performed using forward Euler with *dt* = 0.1.

The model was first constrained by setting *τ*^+^*_s_* = 2 s based on previous literature (Bahl and Engert, 2020; Dragomir, Štih and Portugues, 2020; Slangewal *et al*., 2026). Then, *τ*^+^*_m_* was fixed using the autocorrelation function of the response **(Fig. 3a)**. Here, correlation values slowly decayed with a time constant of around 15 trials (trials are on average 60 s long). We thus set *τ*^+^*_m_* to 900 s. We then tuned the remaining four parameters (*k*^+^*_s_*, *τ*^−^*_s_*, *k*^−^*_s_*, and *k*^+^*_m_*) using a heuristic, controller-tuning procedure analogous to the classical Ziegler–Nichols method (Ziegler and Nichols, 1942): First, *k*^+^*_s_* (weight of the fastest integrator) was set to match the experimentally observed within-trial turn angle peak response (**Fig. 4b** and **Fig. S4g**). Next, we tested several values of *τ*^−^*_s_* (between 10 and 90 s) and varied *k*^−^*_s_* until we could reproduce the repulsion effect (**Fig. 2e**). Finally, we altered *k*^+^*_m_* to reproduce trial-to-trial repetition (**Fig. 2c**). The parameters that were best able to reproduce trial-to-trial effects were selected (*k*^+^*_s_* = 25, *τ*^−^*_s_* = 45 *s*, *k*^−^*_s_* = 15, and *k*^+^*_m_* = 2.4).

## Data and code availability

All experimental and simulation data, Python scripts for data analyses and simulations, as well as to generate plots, are available at the KonData repository with the persistent identifier https://doi.org/10.48606/h5v2506p48g298zv. Requests for further information and resources should be directed to and will be fulfilled by Armin Bahl.

The above-mentioned persistent identifier will only work upon final paper acceptance. For the review process, please use the following temporary link containing the data and source code archives: https://kondata.uni-konstanz.de/radar/en/dataset/h5v2506p48g298zv?token=LPGUwbmZjEoLcLAQQRAo

## Acknowledgments

We would like to thank all the members of the Bahl lab for helpful discussions throughout this work. We thank Ulrike Bonitz and Heike Naumann for administrative support. We thank all the animal caretakers for continuous support in maintaining the fish facility, and our central workshop team for helping build the behavior setups. We also thank Iain Couzin and Vishwesha Guttal for their insights on this project. Furthermore, we thank Roberto Garza, Meha Jadhav, Margherita Zaupa, Daniel Hummel, and Sophie Aimon for reading and commenting on the manuscript draft.

## Author contributions

Conceptualization: A.M., I.H., A.B.; Methodology: A.M., I.H., A.B.; Software: A.M., A.B.; Validation: A.M., I.H., A.B.; Formal analysis: A.M.; Investigation: A.M., S.H.; Resources: A.B.; Data Curation A.M., A.B.; Writing - Original Draft: A.M.; Writing - Review & Editing: A.M., I.H., A.B.; Visualization: A.M.; Supervision: I.H., A.B.; Project administration: A.B.; Funding acquisition: I.H., A.B.

## SUPPLEMENTAL INFORMATION

Document S1

Figures S1 to S5

---

**Figure S1.**
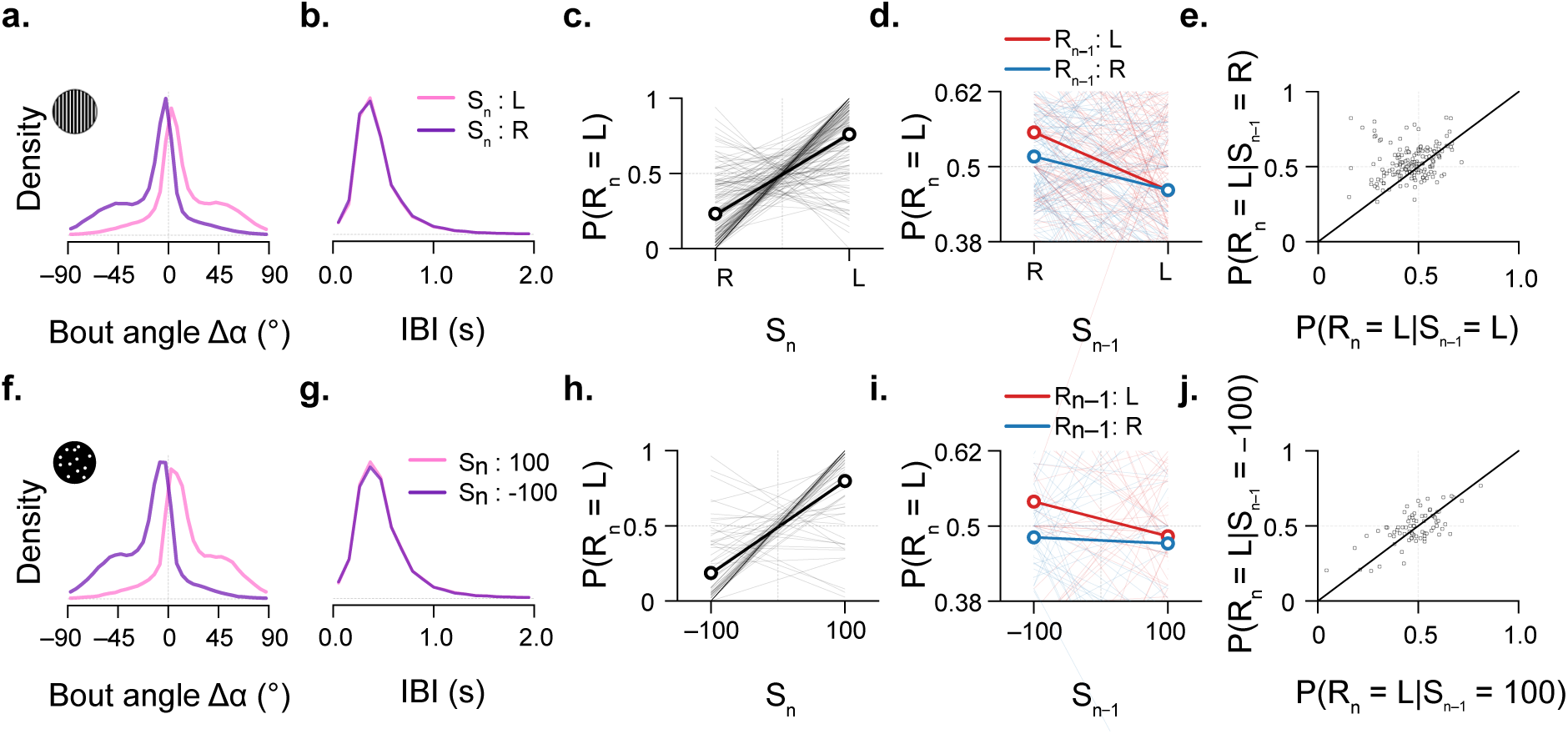
Individual-fish-based analyses also show that larvae modulate responses based on previous trials. **a–e**, Grating experiments. **f–j**, random-dot-motion experiments. **a,f,** Distribution of bout angles. **b,g,** Distribution of interbout intervals (IBI). **c,h**, Probability of a left response to the current stimulus, as in **Fig. 1e,h**, right, but analyzed for individual fish (thin solid lines), rather than through bootstrapping of merged trials. **d,i**, Probability of a left response in the current trial, grouped based on the previous stimulus direction, as in **Fig. 1f,i**, but analyzed for individual fish (thin colored lines). **e,j,** Comparisons of probability of left response in the current trial for different directions of previous stimuli. Each dot is an individual larva. The diagonal is the identity line (y = x). Points falling above the identity line indicate repulsion: fish for which P(left) was higher following *S_n_*_−1_ being rightward motion than following *S_n_*_−1_ being leftward motion (i.e., a history-dependent shift consistent with the population effect). The distribution of fish above vs. below the identity line was tested against the null of an equal split using a two-sided exact binomial (sign) test: 108 of 160 fish above the line, p < 0.001 in (e) and 39 of 76 fish above the line, p = 0.91 in (j). N = 160 fish in (a–e) and N = 76 fish in (f–j). Same animals as in related **Fig. 1**.

**Figure S2.**
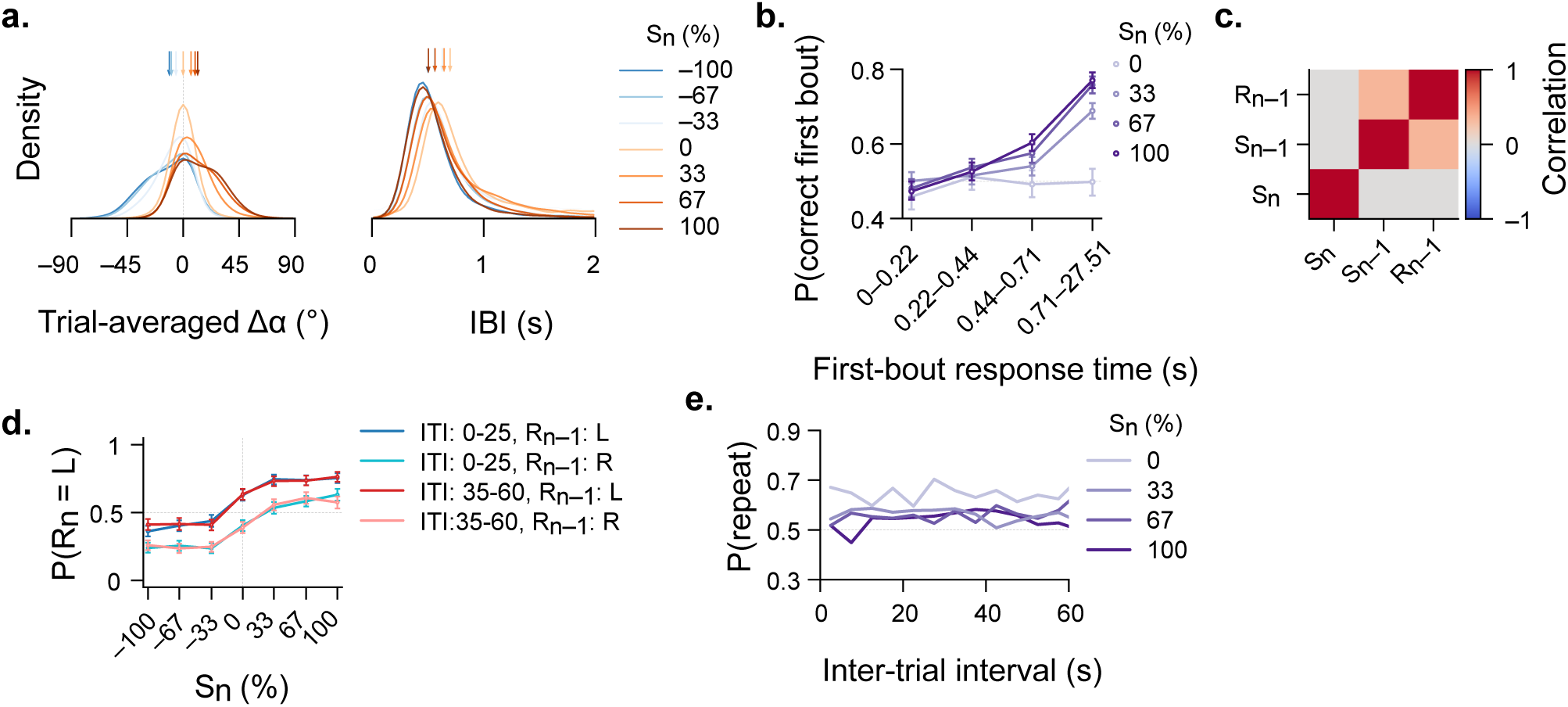
Detailed quantification of behavioral features and history-dependence for the random-dot-motion integration task with variable inter-trial intervals. **a,** Normalized distributions of trial-averaged *Δα* to different coherences (left) and of trial-averaged interbout intervals (IBI) for different coherences (right). Arrows show distribution medians. **b**, Correctness of the first bout after stimulus onset as a function of response time. Monotonic increase test (**Methods**): p < 0.05 for 33%, 67%, and 100% coherence levels. p = 0.37 for 0% coherence level. **c**, Pearson correlation between stimulus and trial-averaged binarized response combinations in the previous and current trials. By design, stimuli are picked randomly (*S_n_* versus *S_n_*_−1_ and *S_n_* versus *R_n_*_−1_). The moderate positive correlation of *S_n_*_−1_ versus *R_n_*_−1_ (0.34), as expected from optomotor behavior, does not compromise the validity of the identified coefficients in our generalized linear model and hierarchical logistic model. **d,** Behavioral performance, grouped on the previous trial response *R_n_*_−1_ direction for short (0 to 25 s) and long (35 to 60 s) inter-trial intervals. Response biases do not seem to depend on interval length. **e**, The probability to repeat the same response for consecutive trials, P(repeat), as a function of inter-trial interval. Also here, history effects are largely independent of time intervals. Same n = 121 larvae as in related **Fig. 2**.

**Figure S3.**
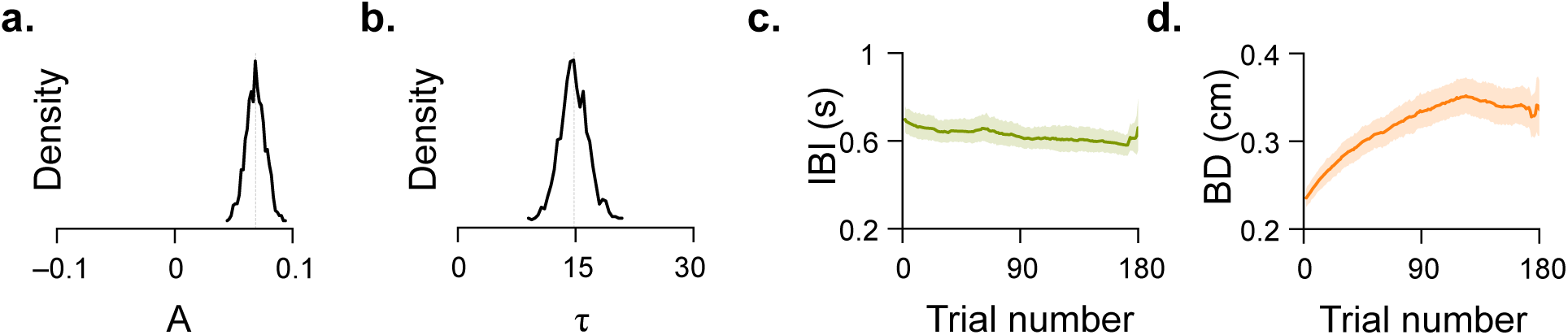
Detailed quantification of the multi-trial autocorrelation and other behavioral metrics over the experiment. **a,b**, Normalized distributions of the autocorrelation value A at lag = 1 trial (a) and exponential decay rate *τ* (b) over 1000 bootstrap iterations. **c**, Interbout interval (IBI) across trials. Spearman correlation between interbout interval and trial was calculated per fish. Mean *ρ* = −0.0672; p < 0.05, Wilcoxon signed-ranked test against *ρ* = 0. **d,** Bout distance (BD) across trials. Same statistics as in (c). Mean *ρ* = 0.2724; p < 0.001. Same N = 121 larvae as in related **Fig. 3**.

**Figure S4.**
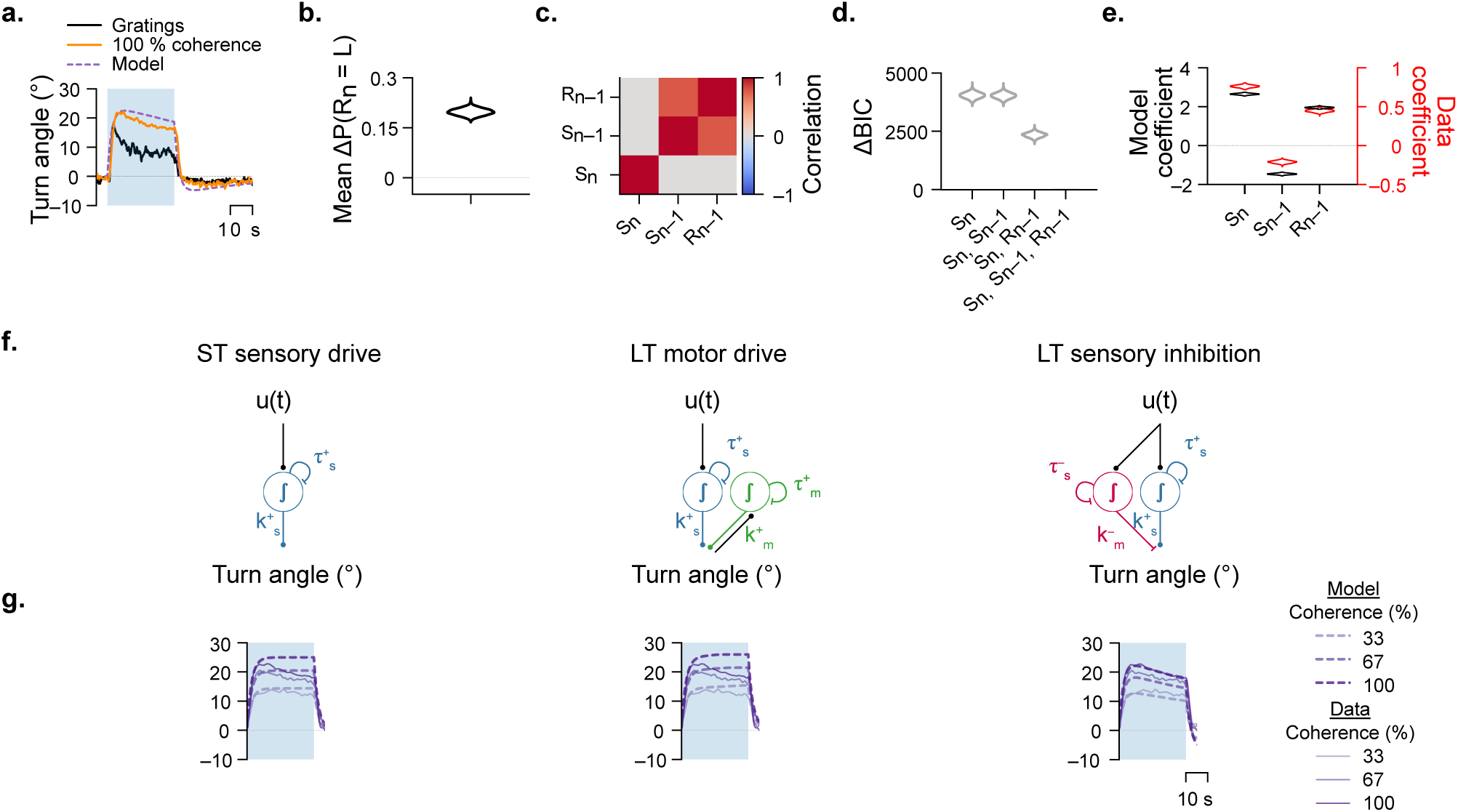
Short and intermediate timescale dynamics and trial-to-trial history in model variants. **a**, Within-trial turn angle dynamics in response to gratings and 100% random-dot-motion stimuli. **b**, Bootstrapped distance of psychometric curves (*R_n_*_−1_ = *L* minus *R_n_*_−1_ = *R*) in **Fig. 4d**. ***p < 0.001 when testing bootstrapped distances against zero (no difference between psychometric curves). **c**, Pearson correlations between *S_n_*, *S_n_*_−1_ with *R_n_*_−1_ from model simulations. The positive correlation of *S_n_*_−1_ versus *R_n_*_−1_ (0.68) does not compromise the validity of the identified coefficients in our generalized linear model. **d,** Comparison of general linear model complexity using ΔBIC. Bootstrapped distance of psychometric curves (*R_n_*_−1_ = *L* minus *R_n_*_−1_ = *R*) in (c). ***p < 0.001 when testing bootstrapped distances against zero (no difference).**e,** Coefficients of predictors in the generalized linear model and data overlaid. **f**, Representation of tested model variants. **g,** Within-trial turn angle dynamics of the alternate models. Model simulations on N = 121 different stimulus streams identical to the ones as in related **Fig. 4**.

**Figure S5.**
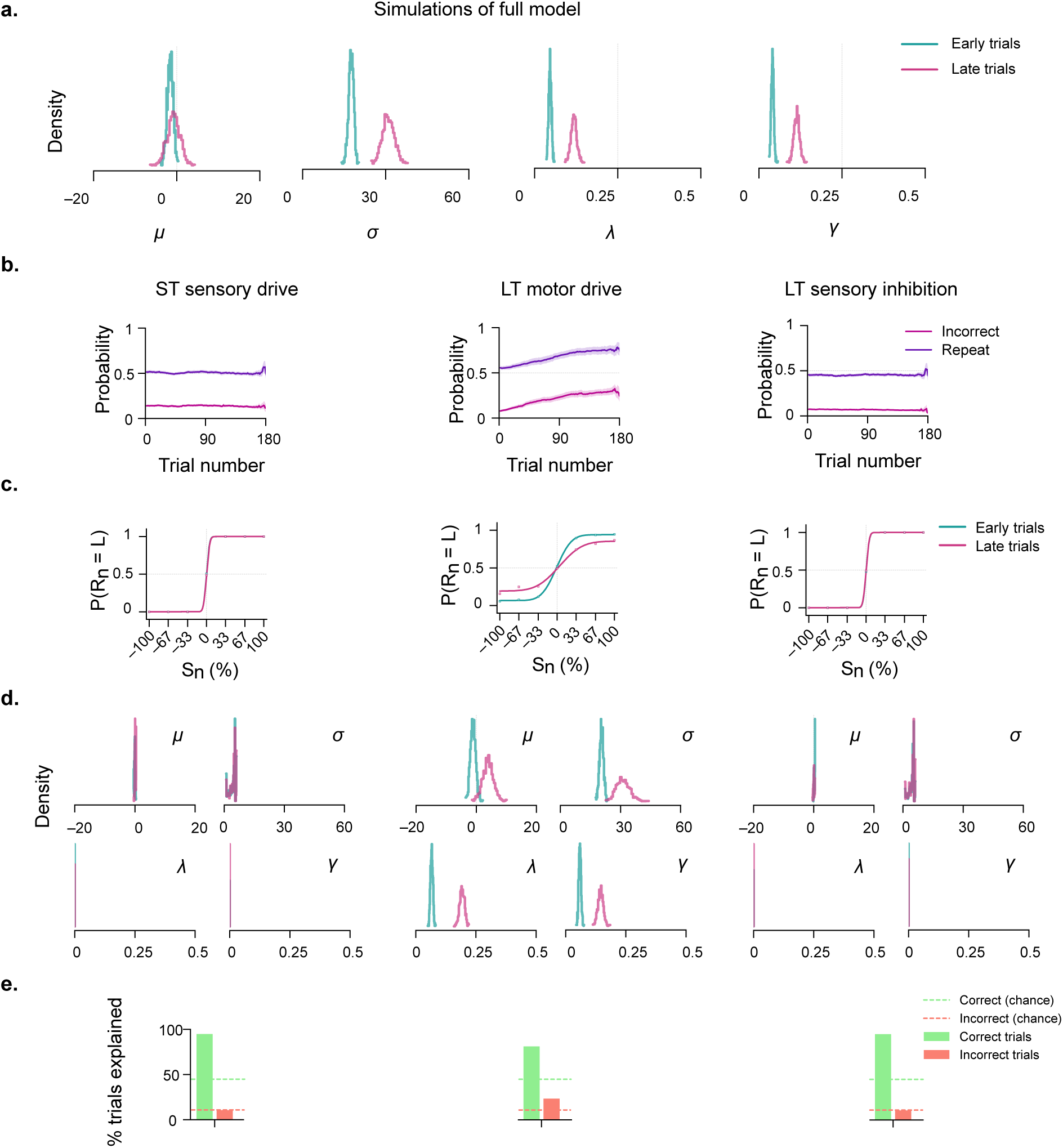
Response repetition and increasing lapse emerge in model simulations. **a,** Normalized distributions of the fitted parameters of the inflection point parameter (*μ*), sensitivity parameter (*σ*), and the lower (*λ*) and upper (*γ*) asymptote parameters for the full model (*Δμ* = 0.799, p=0.652; *Δσ* = 13.428, p <0.001; *Δλ* = 0.072, p < 0.001; *Δγ* = 0.723, p < 0.001, bootstrap test on change in parameters). **b–d,** Quantification of history effects and psychometric curves for the model variants. **b**, Probability to repeat a response and rate of incorrect trials over experimental time. For ST sensory drive: Mean Spearman correlation of repetition over time = 0.005, p = 0.433; Mean Spearman correlation of incorrectness over time = – 0.005, p = 0.46; LT motor drive: Mean Spearman correlation of repetition over time = 0.12, p < 0.001; Mean Spearman correlation of incorrectness over time = 0.143, p < 0.001; LT sensory inhibition: Mean Spearman correlation of repetition over time = 0.001, p = 0.85; Mean Spearman correlation of incorrectness over time = –0.011, p = 0.107; **c,d**, Psychometric curves with fitted parameterization for early (before 70 min) and late (after 120 min) trials. For ST sensory drive : *Δμ* =0.2, p=0.41; *Δσ* = 0.03, p = 0.99; *Δλ* = 0.0, p = 0.69; *Δγ* = 0.0, p = 0.76, bootstrap test on change in parameters; LT motor drive : *Δμ* = 4.96, p=0.01; *Δσ* = 10.7, p < 0.001; *Δλ* = 0.127, p < 0.001; *Δγ* = 0.089, p < 0.001, bootstrap test on change in parameters; LT sensory inhibition: *Δμ* = –0.134, p=0.536; *Δσ* = -0.023, p = 1.00; *Δλ* = 0.0, p = 0.784; *Δγ* = 0.0, p = 0.92. All reported statistics were obtained using bootstrap tests on the change in fitted parameters; this procedure assesses the probability that the observed change includes the null value (*Δ* = 0). **e**, Percentage of correct and incorrect trials above chance explained by the model. Chance levels were computed by shuffling correct and incorrect labels across all trials. Model simulations on N = 121 different stimulus streams identical to the ones used in the experiments. Figure relates to **Fig. 5**.

